# Elucidating the Mechanism by Which HIV-1 Nucleocapsid Mutations Confer Resistance to Integrase Strand Transfer Inhibitors

**DOI:** 10.1101/2025.05.17.654662

**Authors:** Yuta Hikichi, Ryan C. Burdick, Sean C. Patro, Si-Yuan Ding, Brian T. Luke, Erin Clark, Sherimay D. Ablan, Xiaolin Wu, Vinay K. Pathak, Eric O. Freed

## Abstract

Persons with HIV (PWH) receiving integrase (IN) strand transfer inhibitors (INSTIs) have been reported to experience virologic failure (VF) in the absence of resistance mutations in IN. We previously reported that mutations in the viral nucleocapsid (NC) are selected in the presence of the INSTI dolutegravir (DTG) and confer levels of INSTI resistance comparable to those conferred by clinically relevant IN mutations. Here we show that these NC mutations accelerate the kinetics of viral DNA integration. The shortened time frame between the completion of reverse transcription and integration correlates with reduced sensitivity to DTG, suggesting that NC mutations limit the window of opportunity for INSTIs to block viral DNA integration. We find that in primary peripheral blood mononuclear cells, HIV-1 acquires mutations in the viral envelope glycoprotein, NC, and occasionally IN during selection for INSTI resistance. Notably, the selected NC and IN mutations act in concert to reduce the susceptibility of the virus to INSTIs. These results provide insights into the mechanism by which HIV-1 escapes the inhibitory activity of INSTIs and underscore the importance of genotypic analysis outside IN in PWH experiencing VF on INSTI-containing drug regimens.

## Introduction

Combination antiretroviral therapy (ART) has significantly reduced HIV-1-associated morbidity, mortality, and the risk of HIV transmission. Integrase (IN) strand transfer inhibitors (INSTIs) are a key class of antiretroviral with minimal side effects, high potency, and a substantial genetic barrier to resistance compared to other classes of antiretrovirals^1,2^. Since 2018, the World Health Organization (WHO) has recommended the INSTI dolutegravir (DTG) as the preferred first-line and second-line treatment for persons with HIV (PWH). However, with the global implementation of DTG-based regimens, there has been an unexpected increase in virologic failure (VF) cases compared to clinical trial predictions^3^.

Retroviral IN proteins catalyze two enzymatic reactions, 3’ processing and strand transfer. In the first reaction, IN removes two nucleotides from the viral DNA ends to create a 5’ overhang. In the strand transfer reaction, IN creates staggered nicks in the target (host) DNA and inserts the processed viral DNA ends, resulting in integration. IN binds the viral DNA ends as a multimeric complex containing at least four molecules of IN (an IN tetramer). Higher-order IN complexes (e.g., a tetramer of tetramers) may also assemble on the viral DNA ends. The complex of viral DNA ends with an IN multimer is often referred to as the intasome^4,5^. As their name implies, INSTIs block the strand transfer activity of IN by binding the intasome rather than free IN^5–8^. Thus, INSTIs cannot exert their inhibitory activity until the intasome has been assembled and the 3’ ends have been processed. Current FDA-approved INSTIs share two critical pharmacophores; a metal-chelating scaffold that binds two Mg^2+^ cofactors within the IN active site, and a halobenzyl moiety that interacts with the 3’-processed viral DNA ends^9^.

HIV-1 assembly is driven by the Gag precursor protein, which contains several structural and functional domains: matrix (MA), capsid (CA), spacer peptide 1 (SP1), nucleocapsid (NC), SP2, and p6^10^. After virus particles assemble and are released from the cell, the viral protease (PR) cleaves the Gag precursor protein into its mature components – MA, CA, SP1, NC, SP2, and p6 – triggering virus maturation. The NC domain of Gag, and the mature NC protein generated during virion maturation, play a variety of roles both early and late in the viral replication cycle^10,11^. NC contains two Cys-Cys-His-Cys (CCHC) zinc finger-like domains, which are connected by a basic linker domain. During virus assembly, both the basic linker and zinc-finger domains play a central role in the selective packaging of the viral RNA genome into assembling particles. The interaction between viral genomic RNA and the NC domain of Gag also promotes Gag multimerization during assembly ^12^. After virus assembly and maturation, NC binds to and promotes the condensation of the genomic RNA inside the mature core. NC functions as a molecular chaperone, facilitating the remodeling of viral RNA structures to form the most thermodynamically stable conformations for tRNA primer annealing, and initiation, strand transfer, and elongation during reverse transcription ^11,13^. Recent studies suggest that reverse transcription of the flexible viral RNA into the comparatively stiff and inflexible double-stranded DNA copy exerts internal pressures on the core, resulting in uncoating ^14–19^. NC binds to newly synthesized viral DNA and condenses the DNA into a tight globule ^17,20,21^. NC-mediated DNA compaction is hypothesized to reduce the pressure within the viral core, thereby preventing premature uncoating in the cytoplasm. Several studies also support the involvement of NC in viral DNA integration. Early in vitro biochemical studies suggest that NC stimulates IN strand transfer activity, and some zinc-finger domain mutants show enhanced ability over WT in stimulating strand transfer. NC binding to the viral DNA in vitro recruits IN, resulting in the formation of a stable NC-DNA-IN nucleoprotein complex ^22–27^. Mutation of the CCHC zinc finger motifs (specifically the NC-H23C and NC-H44C mutations) results in a virus that is competent in reverse transcription but highly defective in integration ^28–30^. NC has also been reported to form biomolecular condensates together with viral RNA, RT and IN ^31^. Thus, it appears likely that NC plays some role in enhancing viral DNA integration, though the mechanistic details remain ill defined.

In many cases of VF to INSTI-containing regimens, resistance mutations are identified in IN. These mutations often alter the drug-target interaction^32^. However, a growing number of clinical studies, some with verified drug adherence, have documented INSTI VF in the absence resistance mutations in IN ^33–37^. These studies imply that mutations outside the IN-coding region contribute to INSTI resistance. In vitro studies demonstrate that mutations in the 3′ polypurine tract (PPT) confer resistance to INSTIs ^38–44^. These 3’PPT mutations increase the levels of unintegrated 1-LTR circles, which can serve as templates for viral protein expression, particularly in cells expressing HTLV-1 Tax ^41,42^. Witjing et al. reported a distinct set of 3′PPT mutations from an individual failing DTG monotherapy in the absence of IN resistance mutations^44^. However, other studies demonstrated that the in vivo-derived 3′PPT mutations do not confer resistance to INSTIs in vitro ^45,46^. Given the widespread use of INSTIs globally and the approval of long-acting INSTIs in clinical settings, it is crucial to monitor and understand the mechanisms underlying resistance to INSTIs.

We previously reported non-canonical pathways for HIV-1 to acquire high-level resistance to INSTIs. Long-term propagation of HIV-1 in cell culture in the presence of DTG revealed that, independent of viral strain, HIV-1 frequently acquired mutations in the envelope glycoprotein (Env) and NC in the absence of INSTI resistance mutations in IN ^47–49^. The accumulated Env mutations lead to high-level INSTI resistance by increasing the multiplicity of infection (MOI) via enhanced cell-cell transfer efficiency. While changes in Env reduce viral susceptibility to multiple classes of antiretrovirals, the fold resistance was notably higher for INSTIs than for other classes of antiretrovirals. Further analysis revealed that the ability of INSTIs to block virus infection is more readily overwhelmed by high MOI compared to other drug classes^49^. We also demonstrated that mutations in NC reduce sensitivity of HIV-1 to INSTIs but not to other classes of antiretrovirals, and NC mutations were not identified during propagation of the virus in the presence of an RT inhibitor. These findings indicated that resistance conferred by NC mutations is specific to INSTIs; however, the mechanism by which NC mutations reduce the susceptibility of HIV-1 to INSTIs was not investigated^49^.

In the present study, we show that the NC mutations selected in the presence of DTG do not have major effects on virus assembly, infectivity, or replication. However, we demonstrate that they reduce the time between completion of reverse transcription and integration, thereby limiting the amount of time during which INSTIs can effectively bind to the intasome and inhibit integration. We also show that NC mutations act in concert with IN mutations to increase INSTI resistance. Our findings provide new insights into the role of NC in the early stages of HIV-1 replication and reveal a new mechanism of INSTI resistance. These findings also support the need for genotypic analysis outside of IN in PWH failing INSTI-containing regimens.

## Results

### NC mutations confer resistance to DTG in T cells without apparent fitness costs

Our previous study demonstrated that HIV-1 frequently acquires NC mutations in the presence of DTG independent of viral strain^49^. These selected NC mutations reduce sensitivity to INSTIs but not to other classes of antiretrovirals, indicating that the resistance conferred by NC mutations is specific to INSTIs. Here we first examined the replication of the NC mutants in the SupT1 T-cell line, previously used for the DTG resistance selection experiments in cell culture ^49^. As controls, we included NC-H44C, an NC-CCHC motif mutant^29^; IN-D116N, a catalytically inactive IN mutant^50^; and 3’PPT-2C3A5T6Δ^38,39^. Some 3’PPT mutants are reported to confer high-level resistance to INSTIs due to integration-independent viral replication under specific conditions; e.g., in cell lines expressing HTLV-1 Tax ^39,41,42^. NC-G19S (**Fig. 1A**) and most other selected NC mutants (**Extended data 1**) exhibited WT replication kinetics in the SupT1 T-cell line in the absence of DTG, indicating that the NC mutations imposed minimal fitness cost. In contrast, the control mutants NC-H44C, IN-D116N, and 3’PPT-2C3A5T6Δ failed to replicate in the SupT1 T-cell line. We further examined the replication in the presence of varying concentrations of DTG (**Fig. 1A and Extended data 1**). At 1 – 3 nM DTG, the NC mutants, except for NC-N27K and R29G, demonstrated a replication advantage over WT NL4-3. By calculating IC_50_ based on the area under the curve (AUC) of the replication kinetics from three repeat experiments, we determined that the NC mutations confer ∼2 to 3-fold resistance to DTG in the SupT1 T-cell line in the context of multi-cycle replication (**Fig. 1B**). As an additional approach to measuring the level of DTG resistance conferred by the NC mutations in the SupT1 T-cell line, we performed single-cycle DTG sensitivity assays using a VSV-G-pseudotyped HIV-1 nanoLuc reporter virus ^51,52^ (**Fig. 1C and D**). Consistent with the multi-cycle replication data, the selected NC mutants, except for NC-R29G, conferred ∼2.8 to 8-fold resistance to DTG. To assess DTG resistance in a more physiologically relevant cell type, we examined DTG sensitivity of the NC mutants in primary CD4+ T cells (**Fig. 1E and F**). Although donor-dependent variability was observed, all of the NC mutants increased DTG IC_50_ in primary CD4 T cells by ∼2 to 6-fold. Together, these data indicate that the selected NC mutations reduce the susceptibility of HIV-1 to DTG, irrespective of target cell type.

**Fig. 1.**
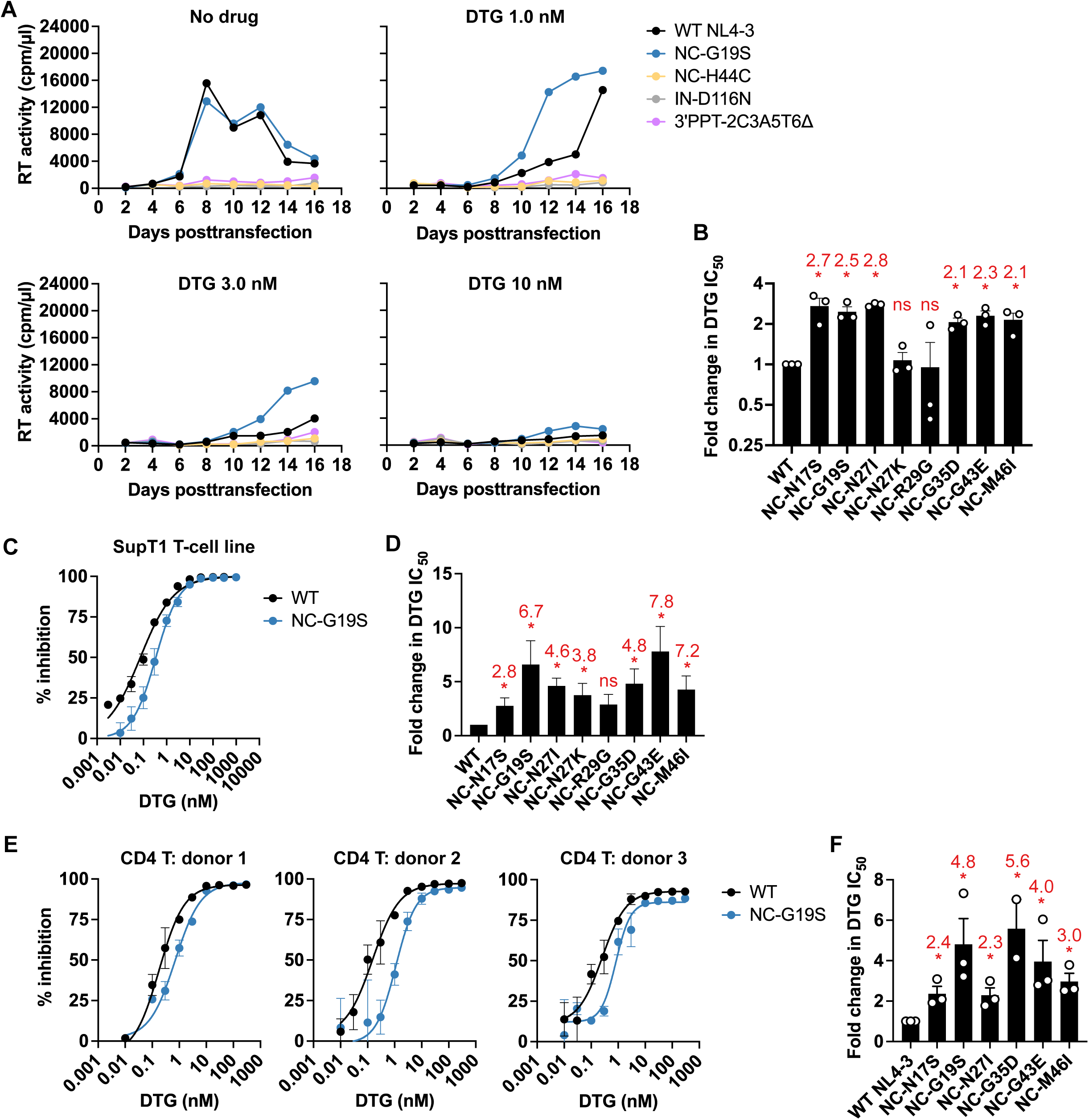
NC mutations reduce the susceptibility of HIV-1 to DTG in T cells. (A) Replication kinetics of the indicated NL4-3 variants in the SupT1 T-cell line in the absence or presence of DTG. Replication curves obtained in the presence of 0, 1, 3, and 10 nM DTG are shown. Data are representative of three independent experiments. (B) Fold changes in IC_50_ were calculated compared to that for the WT over a range of DTG concentrations (0.01 to 300 nM). IC_50_ values were calculated on the basis of the area under the curve (AUC) of the replication kinetics. Fold changes in IC_50_ are shown above the graph. \**P* < 0.05, one-sample *t* test. (C - D) DTG sensitivity of the NC mutants in the SupT1 T-cell line. SupT1 T-cells were incubated with RT-normalized VSV-G-pseudotyped nanoLuc reporter HIV-1 in the presence of various concentrations of DTG. Nano luciferase activity was measured at 48 h post-infection. Data from at least three independent experiments are shown as means ± SEM. Fold changes in IC_50_ are shown in D. \**P* < 0.05, one-sample *t* test. (E) DTG sensitivity of the NC mutants in primary human CD4^+^ T cells. Primary CD4^+^ T cells from the three healthy donors were incubated with RT-normalized VSV-G-pseudotyped nanoLuc reporter HIV-1 in the presence of varying concentrations of DTG. Nano luciferase activity was measured at 48 h post-infection. Error bars indicate the standard error of the duplicate infection. (F) Fold changes in IC_50_ based on the data in panel E. Data are shown as means ± SEM. \**P* < 0.05, unpaired *t* test.

### NC mutations selected in the presence of DTG do not disrupt virus assembly or infectivity

We previously demonstrated that the NC mutations selected in the presence of DTG, most of which cluster in the zinc-finger domains (**Extended data 2A)**, do not affect single-cycle, cell-free infectivity^49^. We next examined whether the NC mutations affect Gag processing or viral release efficiency by western blotting (**Extended data 2B and C**). The analysis indicated that the selected NC mutations in the zinc-finger domain do not affect Gag processing in virus-producing cells or in virions (**Extended data 2B-E**), virus release efficiency (**Extended data 2F**), or the amount of virion-associated reverse transcriptase (RT) protein (p66/p51) (**Extended data 2G**). Several mutations in the basic linker (R29M, R32G, and G35N) slightly decreased (by <two-fold) virus release efficiency (**Extended data 2F**). The zinc-finger domains are crucial for nucleic acid binding and RNA packaging into the virion^12^. Specifically, previous reports indicated that mutations in the NC zinc-finger CCHC motifs (e.g., NC-H23C and H44C) impair viral RNA packaging into the virion and reduce particle infectivity^30,53,54^. To investigate the effect of the NC mutations selected in the presence of DTG on RNA packaging, we measured the amount of viral RNA in WT and mutant virions by real-time PCR (**Extended data 2H**). We found that the NC-N17S and G35N mutations slightly decrease viral RNA packaging; however, most of the selected NC mutations do not exhibit altered viral RNA packaging. As controls, we tested NC-H23C and H44C mutants and confirmed previous reports^30,53,54^ that they markedly reduce RNA packaging (**Extended data 2H**). These results indicate that the selected NC mutations have little or no effect on virus assembly or particle infectivity.

### NC mutations selected in the presence of DTG do not alter the viral DNA ends

Previous studies indicate that INSTIs selectively interact with the intasome, a complex of multimeric IN and the viral DNA ends, but do not bind tightly to free IN^5–8^. Consequently, modification of the viral DNA ends could disrupt the INSTI-intasome interaction^55,56^, potentially leading to INSTI resistance. However, modifications of the viral DNA ends would be expected to cause severe defects in infectivity ^57,58^, which, as shown above, is not observed for the selected NC mutants. Newly formed viral DNA ends can either serve as substrates for integration into the host genome – the reaction catalyzed by IN – or can undergo a non-homologous end-joining reaction in the nucleus to form 1-LTR and 2-LTR circular forms^59^. These circular DNAs are not substrates for integration but sequencing across the 2-LTR junctions provides information about the integrity of the viral DNA ends. In the presence of the catalytically inactive IN-D116N mutation, integration does not occur and circular DNA forms accumulate in the nucleus^29,50^. To investigate whether the NC mutations selected in the presence of DTG alter the integrity of the viral DNA ends, we sequenced the 2-LTR junctions of circular viral DNA from cells infected with WT and three representative NC mutants selected in the presence of DTG: NC-N17S, G19S, and N27I. We mainly observed four patterns of 2-LTR junctions in the WT-infected cells: an intact GTAC motif at the junction, small modifications in the GTAC motif, large deletions in the LTR regions, and insertions at the junction (**Extended data 3A - B**). The NC-N17S, G19S, and N27I NC mutations did not significantly alter the distribution of 2-LTR junction sequences compared to WT (**Fig. 2A – B, Extendend data 3C**). As controls in these experiments, we included NC-H44C^29^ and 3’PPT-2C3A5T6Δ^38,39^. Previous studies demonstrated that NC-H44C cannot create the viral DNA ends necessary for integration^29,30^. Consistent with these previous reports, we observed that, compared to WT, the NC-H44C mutant exhibited a lower frequency of consensus 2-LTR junctions, and a higher frequency of 2-LTR junctions appended with long insertions or substitutions (**Fig. 2A and Extended data 3C**). It has been shown that modification of the 3’PPT G-tract abrogates RNase H cleavage during reverse transcription, leading to the retention of the 3’PPT in the U3 region^60–62^. Consistent with these studies, DTG-resistant 3’PPT mutations caused the formation of 2-LTR junctions with an extended U3 region due to the retention of 3’PPT sequences (**Fig. 2A and Extended data 3A and C**). This specific modification at the junction was not detected in cells infected with the selected NC variants (**Fig. 2A**). These results suggest that the NC mutations confer INSTI resistance via a mechanism distinct from that of the 3′PPT mutant. Indeed, the reported 3’PPT mutations selected with DTG significantly reduce both infectivity and the maximum inhibitory effects of INSTIs^39^, likely due to viral gene expression from unintegrated episomal DNA ^40,41^. In contrast, NC-G19S does not affect infectivity but causes a ∼3-fold increase in the DTG IC_50_ (**Extended data 4A - B**).

**Fig. 2.**
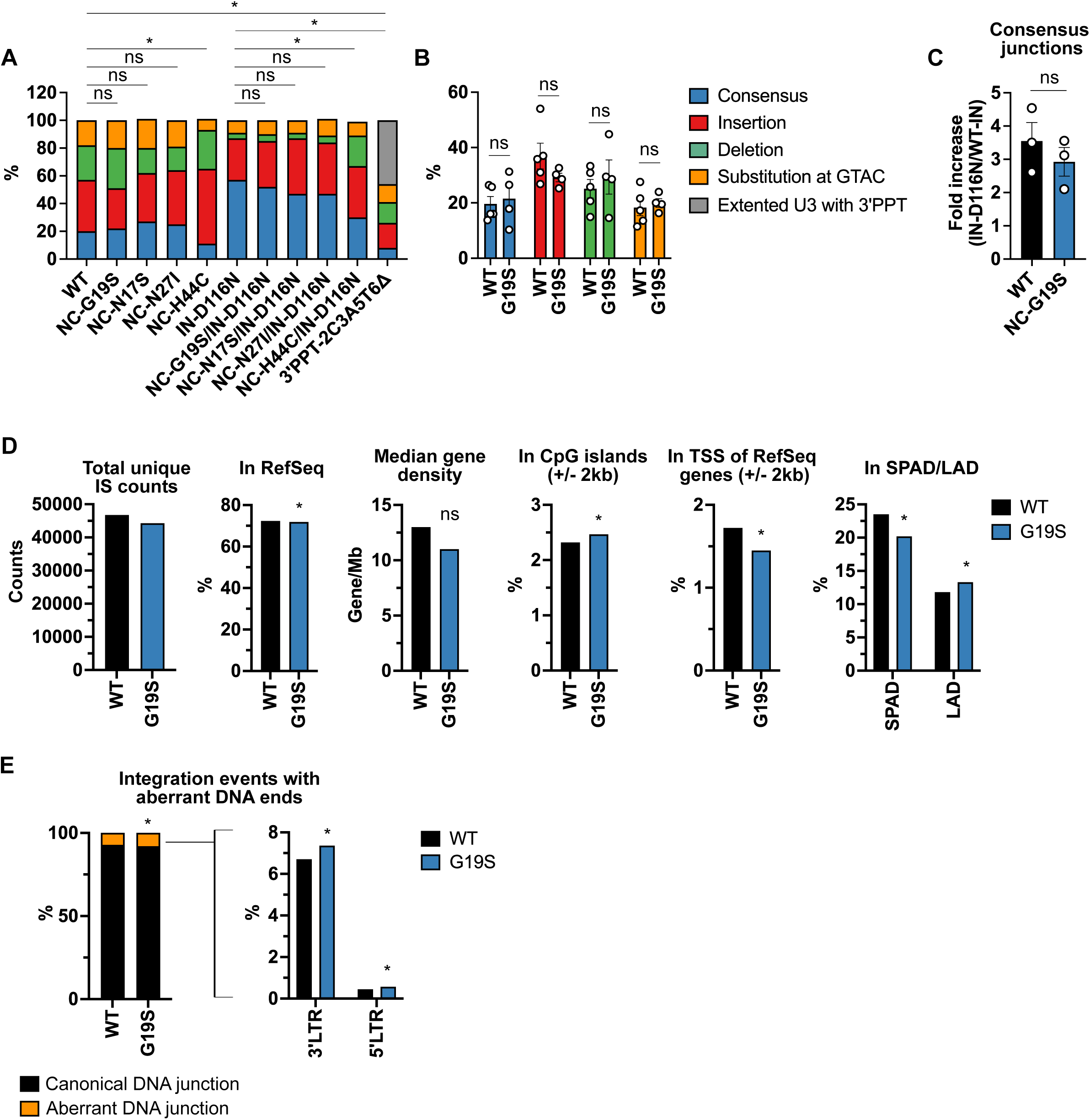
Sequencing analysis of NC-mutant 2-LTR junctions. (A) Frequency distribution of 2-LTR junction categories (consensus GTAC, insertion, deletion, subsitutions in GTAC, extended U3 with 3’PPT) in cells infected with WT HIV-1 or the indicated mutants. The chi-squared test was used to compare overall trends in modified 2-LTR junction frequencies (*P < 0.05). ns: not significant) (B) Comparison of specific 2-LTR junction categories between WT and NC-G19S mutant. Data represent means ± SEM from 4 - 5 independent experiments. Statistical significance was determined using unpaired t-test (*p < 0.05). ns: not significant. (C) Fold increase in the consensus sequence frequency at 2-LTR junctions in IN-D116N variants compared to WT-IN variants. Statistical significance was evaluated using an unpaired t-test (*p < 0.05). ns: not significant. (D) Integration site (IS) frequency in cells infected with WT or NC-G19S. The total number of unique ISs, gene density within a 1 Mb region surrounding each IS, and the frequency of integration within RefSeq genes (genes annotated in the NCBI Reference Sequence Database), CpG islands (±2 kb), transcription start sites (TSS) of RefSeq genes, speckle-associated domains (SPADs), and lamina-associated domains (LADs) are shown. The binomial test was used to compare WT and NC-G19S (*p < 0.05). ns: not significant. (E) The frequency of the integration events with canonical and aberrant viral DNA ends at 5’LTR and 3’LTR. The binomial test was used to compare WT and NC-G19S (*p < 0.05). ns: not significant.

Because the IN-D116N mutant is catalytically inactive and unable to direct 3’ processing, it exhibits an increase in the prevalence of unprocessed consensus GTAC motifs at the 2-LTR junction compared to WT ^29,50^. To further investigate whether the selected NC mutations affect the viral DNA ends, we introduced the IN-D116N mutation into WT and NC-mutant molecular clones. As reported^29^, IN-D116N exhibited an increase in consensus GTAC sequences at the 2-LTR junctions (**Fig. 2A**). The selected NC mutants in the context of IN-D116N displayed a similar distribution of 2-LTR junction sequences as IN-D116N (**Fig. 2A and Extended data 3C**). The control mutant, NC-H44C/IN-D116N, exhibited a lower frequency of consensus junctions and a higher frequency of deletions compared to IN-D116N (**Fig. 2A and Extended data 3C**). Notably, the fold increase in the consensus junction measured with IN-D116N is not altered by NC-G19S, suggesting that the NC mutations do not disrupt 3’processing events before integration (**Fig. 2C**).

To investigate in more detail the impact of the selected NC mutations on formation of viral DNA ends, we performed integration site sequencing coupled with bioinformatic analysis^63^. For this purpose, we used the NC-G19S mutant. HIV-1 preferentially integrates its viral DNA in transcriptionally active, gene-dense regions known as speckle-associated domains (SPADs), and disfavors integration in heterochromatin regions such as lamina-associated domains (LADs) ^64^. Several host factors, in particular cleavage and polyadenylation factor 6 (CPSF6), play key roles in HIV-1 integration site targeting^64^. This analysis indicated that NC-G19S exhibited a slight decrease in the frequency of integration sites in SPADs and a corresponding increase in integration into LADs (**Fig. 2D and Extended data 3D**). However, the frequency of integration events with aberrant viral DNA ends is rare (**Fig. 2E and Extended data 3E**). Specifically, with ∼50,000 integration junctions sequenced, WT and NC-G19S showed 92.84% and 92.07% canonical DNA ends, respectively. Of the 7.16% of ends that were aberrant for WT, 0.45% were at the 5’ LTR and 6.71% were at the 3’ LTR. Of the 7.93% of ends that were aberrant for NC-G19S, 0.57% were at the 5’ LTR and 7.36% were at the 3’ LTR (**Fig. 2E and Extended data 3E**). While these differences were statistically significant, likely due to the large sample size, their biological relevance is uncertain. These results are consistent with the observation that the selected NC mutations exhibit infectivity and replication capacity comparable to that of the WT^49^ (**Fig. 1 and Extended data 1 and 4A**). Taken together, we conclude that the NC mutations selected with DTG do not decrease the sensitivity of the virus to INSTIs by compromising the integrity of the viral DNA ends.

The WT levels of infectivity measured with NC-G19S and the other NC mutants obtained from the selections in DTG suggest that these NC mutants are able to form functional intasomes. To delve more deeply into this question, we used an orthogonal approach. We previously reported that the antiviral activity of INSTIs can be readily overwhelmed by a high multiplicity of infection (MOI) ^49^. We developed an abrogation assay based on co-infecting target cells with a fixed amount of VSV-G-pseudotyped GFP reporter virus with varying amounts of VSV-G-pseudotyped virus lacking a fluorescent reporter (referred to as "dark virus") (**Fig. 3A**). By using this assay, we demonstrated that the dark virus abrogated INSTI inhibition of the GFP reporter virus in a manner dependent on dark virus input ^49^ (**Fig. 3B**). In contrast, dark viruses harboring inactivating mutations in RT did not abrogate ^49^ because they do not synthesize viral DNA, and the IN-D116N mutant did not abrogate because it does not carry out the 3’processing step required to form an intasome^4^. If NC-G19S were impaired in its ability to form a functional intasome, we would expect that the NC-G19S dark virus would not rescue the GFP reporter virus from INSTI inhibition; i.e., it would not abrogate the inhibitory activity of the INSTI. As shown in **Fig. 3B-D**, both WT and NC-G19S dark virus are similarly able to rescue GFP expression, in a manner proportional to virus input. As additional controls performed in parallel, we also evaluated the intasome-deficient mutants (IN-D116N and NC-H44C) and the integration-independent 3’PPT mutant (**Fig. 3C – D**). As expected, the intasome-deficient and the 3’PPT mutants failed to rescue GFP expression. These data are consistent with the infectivity assays and demonstrate that NC-G19S forms functional intasomes with an efficiency comparable to that of WT.

**Fig. 3.**
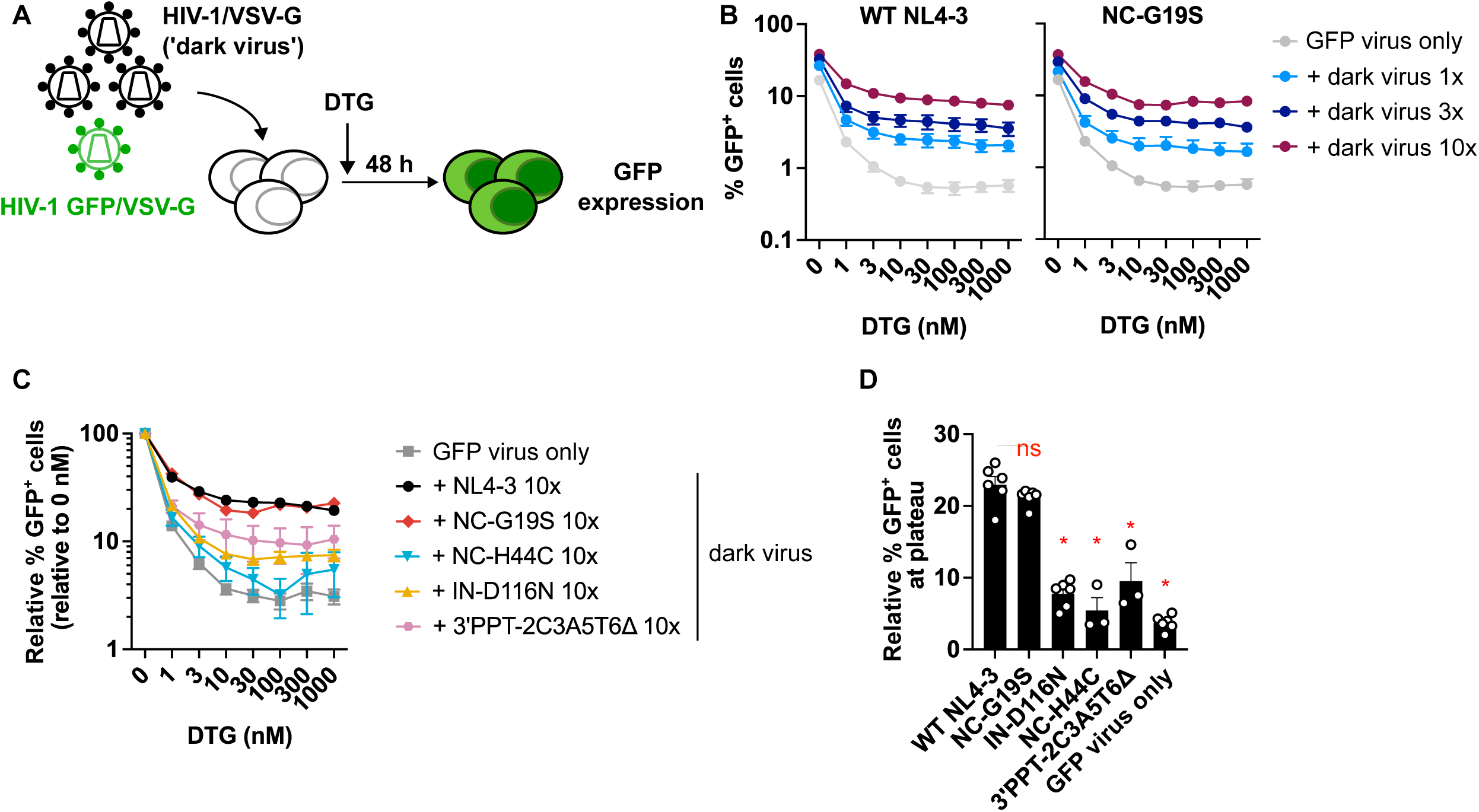
The NC-G19S mutant is similar to WT in its ability to abrogate INSTI inhibition. (A) Experimental scheme for abrogation assay whereby high MOI of non-reporter HIV-1 can abrogate the ability of DTG to block infection of a GFP-reporter HIV-1. B) The SupT1 T-cell line was co-infected with Env(-) VSV-G–pseudotyped HIV-1 (‘dark virus’) at a range of inputs along with a fixed amount of Env(-) VSV-G–pseudotyped GFP reporter virus encoding either WT NC or the NC-G19S mutant. The dark virus input was 1, 3, or 10 times the amount of the GFP reporter virus, as normalized by RT activity. Data are shown as the relative percentage of GFP^+^ cells, normalized to the GFP signals in the absence of DTG. (C) Dose-response curve of DTG against a fixed amount of Env(-) VSV-G–pseudotyped GFP reporter virus, co-infected with 10 times more dark virus harboring the indicated mutations. (D) Summary of the relative percentage of GFP^+^ cells at plateau levels when co-infected with a 10-fold excess of dark virus, as normalized by RT activity. Data are presented as means ± SEM from >3 independent experiments, with statistical significance indicated (*p < 0.05) as determined by one-way ANOVA with Dunnett’s test, ns: not significant.

### The NC-G19S mutation selected in the presence of DTG accelerates early post-entry events including viral DNA integration

NC plays a key role in post-entry events during viral infection ^11,13^ (see Introduction). We previously reported that the NC mutants selected in the presence of DTG exhibit infectivity comparable to that of WT 48 h post-infection in the TZM-bl indicator cell line^49^. To examine the kinetics of early infection events, we measured luciferase activity in TZM-bl cells at both 24 and 48 h post-infection. We found that several of the NC mutants selected in the presence of DTG exhibit significantly higher luciferase signals than the WT at 24 h but not at 48 h (**Fig. 4A**). Importantly, the infectivity of the NC mutants at 24 h but not 48 h post-infection correlates with their level of DTG sensitivity (**Fig. 4B**); as a general trend, the higher the luciferase expression at 24 h the higher the DTG IC_50_. This is consistent with the hypothesis that INSTI resistance conferred by the NC mutations is positively correlated with the kinetics of early post-entry events.

**Fig. 4.**
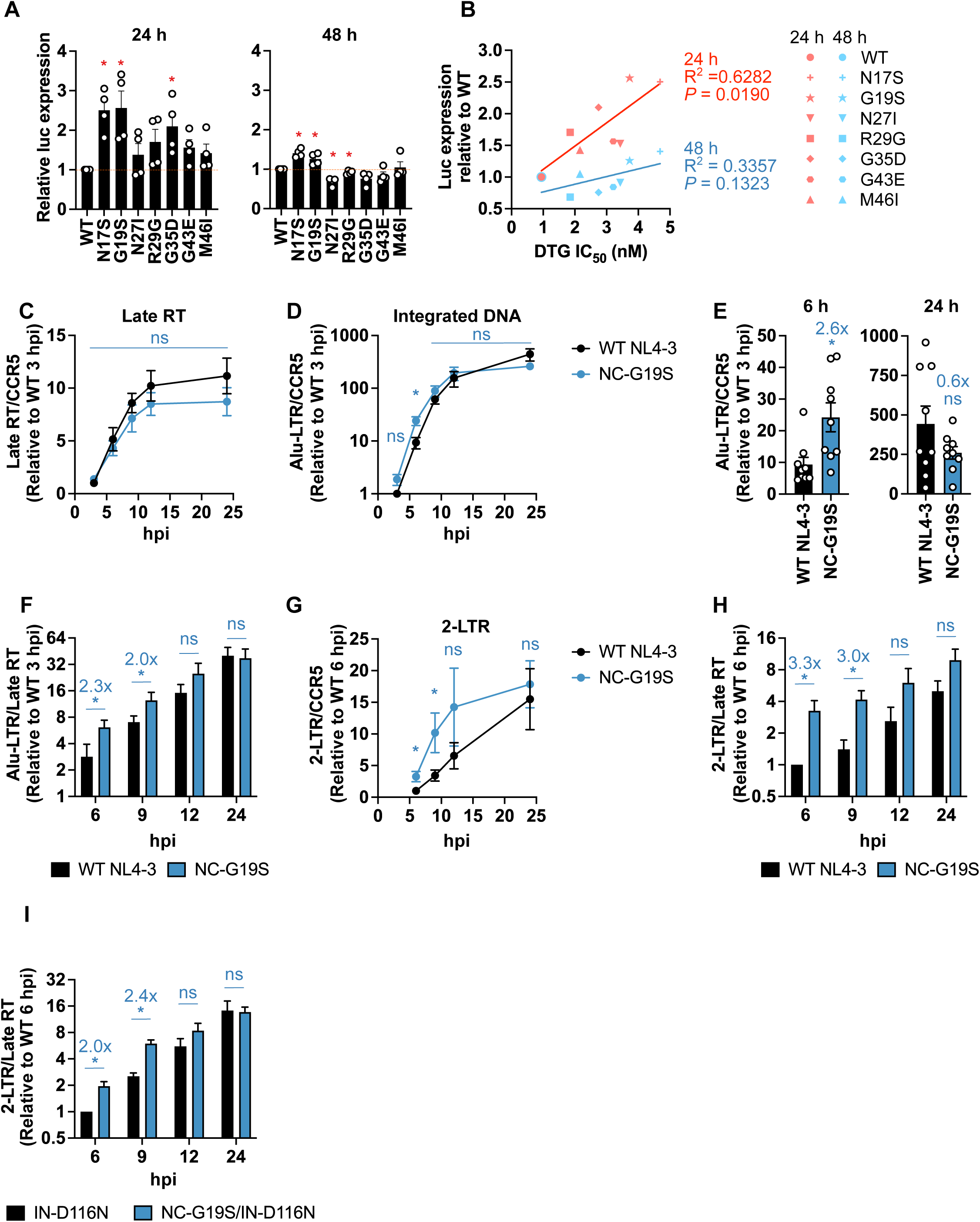
The NC-G19S mutant accelerates 2-LTR circle formation and viral DNA integration. (A) RT-normalized WT and NC-mutant virus stocks were used to infect TZM-bl cells. Relative infectivity of the indicated NC mutants compared to WT at 24 and 48 h is shown. Statistical significance is shown (*p < 0.05) as determined by a one-sample t-test. (B) Correlation between the relative infectivity at the indicated time points and the DTG IC_50_. DTG IC_50_ was measured at 48 h post-infection (Mean IC_50_ values are from Hikichi et al., Sci Adv., 2024^49^). SupT1 T-cells were exposed to Env(-), DNase I-treated, VSV-G-pseudotyped WT or NC-G19S virus particles and the indicated viral DNA species were quantified by qPCR. The cellular DNA was extracted at the indicated time points and subjected to qPCR using U5/gag primers for (C) late reverse transcription products (late RT), and (D - E) Alu-LTR primers for integrated DNA. (F) The ratio of integrated DNA to the total late RT product at the indicated time point. (G) The kinetics of 2-LTR circle formation. (H – I) The ratio of 2-LTR circles to the total late RT product at the indicated time point of (H) IN-WT and (I) IN-deficient IN-D116N viruses. The data are shown as means ± SEM from at least three independent experiments with statistical significance indicated (*p < 0.05) as per unpaired t-test, ns: not significant.

To investigate which early step(s) in the virus replication cycle are accelerated by the NC mutants, we analyzed the impact of the prototype NC-G19S mutation on the kinetics of reverse transcription, 2-LTR circle formation, and integration by qPCR. The NC-G19S mutation did not significantly alter the kinetics of reverse transcription in the infected SupT1 T-cell line (**Fig. 4C**). However, a nested Alu-LTR qPCR assay revealed that this mutant exhibited approximately 2.6-fold more integrated DNA than WT at 6 h post-infection, although the WT and NC-G19S mutant generated similar levels of integrated DNA at later time points (**Fig. 4D and E**). Notably, the NC-G19S displayed a higher ratio of integrated DNA to total late RT product at 6 - 9 h post-infection, suggesting that NC-G19S undergoes more rapid integration compared to WT (**Fig. 4F**). Next, we quantified the formation of 2-LTR circles, a surrogate marker for viral DNA in the nucleus. The NC-G19S exhibited more rapid accumulation of 2-LTR circles compared to the WT, consistent with the trends of integrated DNA (**Fig. 4G - H**). Previous reports have demonstrated that defects in integration lead to the accumulation of 2-LTR circles in the nucleus^50,65^. To further elucidate the proportion of viral DNA in the nucleus, we quantified 2-LTR circles for the NC-G19S mutant in the context of the IN-D116N catalytic site IN mutant. Consistent with the data obtained with WT IN (**Fig. 4H**), in the context of IN-D116N, the NC-G19S mutant showed similar kinetics of reverse transcription but more rapid accumulation of 2-LTR circles compared to IN-D116N with WT NC (**Fig. 4I**). These data indicate that the NC-G19S mutation accelerates early post-entry events, particularly following the completion of reverse transcription.

To gain more insights into the relative timing of post-entry events, we performed time of drug addition (TOA) assays, which measure the time required by the virus to lose susceptibility to a particular inhibitor. We used the entry inhibitor BMS-806; the RT inhibitor rilpivirine (RPV); the capsid inhibitor lenacapavir (LEN), which is known to disrupt capsid function^19,66,67^; and the INSTI DTG. As reported ^19,68^, WT NL4-3 exhibited time lags between the loss of inhibition by each inhibitor (**Fig. 5A and B**), representing the sequential completion of entry, reverse transcription, uncoating, and integration. WT and NC-G19S exhibited loss of inhibition to BMS-806, RPV, and LEN with similar kinetics (**Fig. 5B - C**). However, compared to WT, NC-G19S lost sensitivity to DTG significantly faster than WT. These TOA results indicate a shorter time window between the completion of reverse transcription and integration, and between uncoating and integration (**Fig. 5D**). These results were reinforced by analyzing the other NC mutants in our panel. Like NC-G19S, these mutants showed a shorter time lag between the completion of reverse transcription/uncoating and integration (**Fig. 5C and D**). These data demonstrate that the NC mutants integrate their viral DNA more rapidly after the completion of reverse transcription and uncoating compared to the WT. To validate these observations with a different assay system, we performed TOA assays using HeLa cells and a VSV-G pseudotyped HIV-1 encoding an mScarlet fluorescent reporter (**Extended data 5**). Consistent with the data presented in **Fig. 5A-D**, NC-G19S did not show a statistically significant difference in the time to loss of sensitivity to the RT inhibitor RPV or the capsid inhibitor LEN but showed an increase in the kinetics of viral DNA integration, resulting in a shorter time between completion of reverse transcription and integration.

**Fig. 5.**
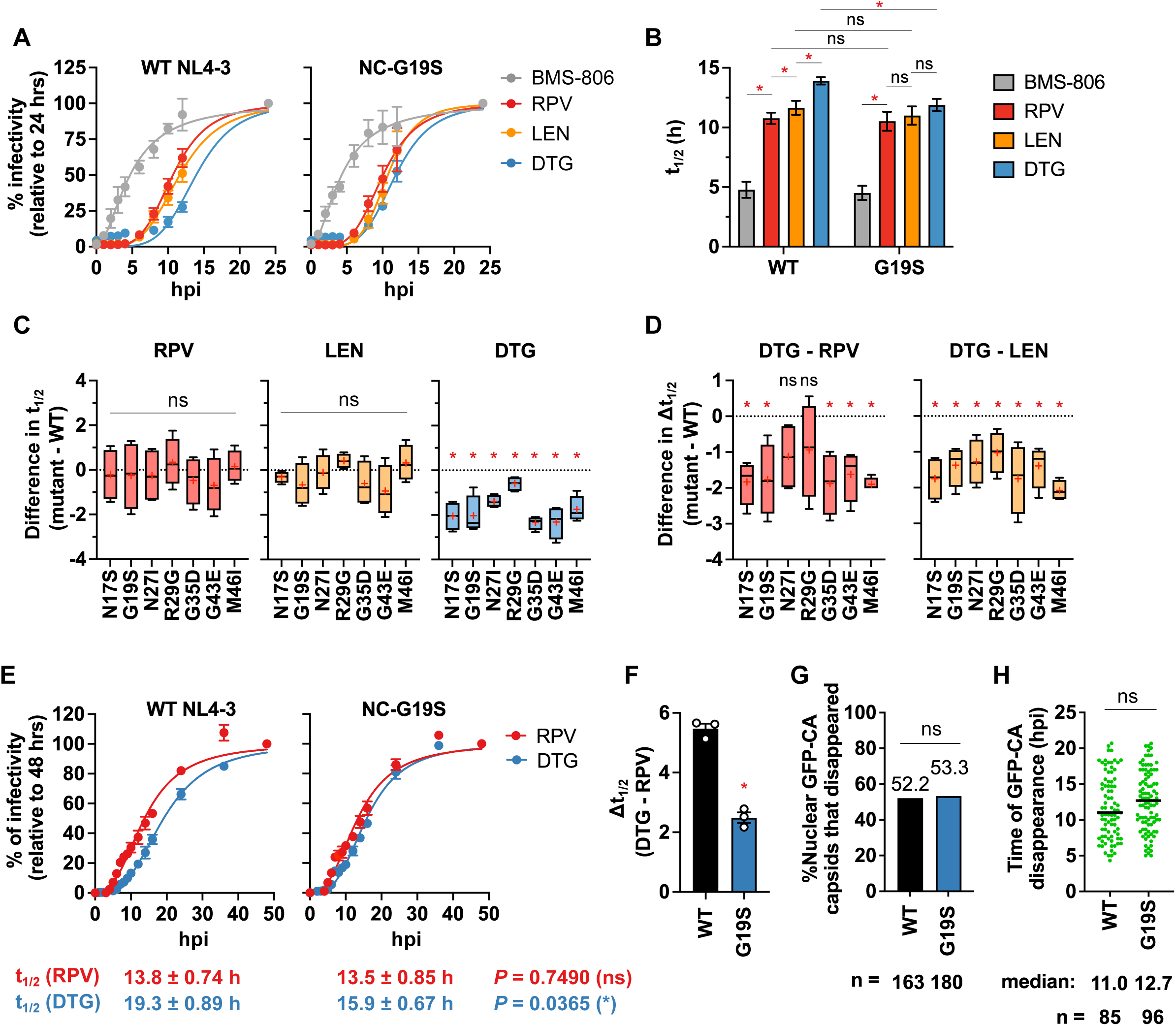
NC mutants shorten the time between completion of reverse transcription and viral DNA integration. (A-B) Time-of-drug addition assay using TZM-bl cells. Drugs were added at the indicated hours post-infection (hpi). Luciferase signals were normalized to the signals at 24 h post-infection. Time to 50% loss of inhibition (t_1/2_) for the indicated drugs is shown in (B). The data are shown as means ± SEM from more than three independent experiments with statistical significance indicated (*p < 0.05) as per mixed-effects model with the two-stage step-up method of Benjamini, Krieger and Yekutieli, ns: not significant. (C) Differences in t_1/2_ for the indicated drugs between WT and the indicated NC mutants. (D) Differences in Δt_1/2_ (the difference in t_1/2_ between DTG and other drugs) between WT and the indicated NC mutants. Box plots show median, interquartile range, minimum, and maximum values. The mean value is indicated by a ‘+’. Statistical significance is shown (*p < 0.05) as determined by a one-sample t-test, ns: not significant. (E) Time of addition assay in SupT1 T-cell line. SupT1 cells were infected with VSV-G-pseudotyped mScarlet reporter virus, followed by the addition of RPV and DTG at the indicated time points. mScarlet expression at the indicated time points was normalized to the signals at 48 h post-infection. t_1/2_ for the indicated drugs is shown below the graph. (F) The difference between t_1/2_ (RPV) and t_1/2_ (DTG). The data are shown as means ± SEM from more than three independent experiments with statistical significance indicated (*p < 0.05) as per unpaired t-test. (G) Percentage of nuclear HIV-1 cores that disappeared during live-cell imaging for the indicated virus in HeLa cells. (H) Time after infection until disappearance of GFP-CA in the nucleus. The total particle counts from three independent experiments are shown below the graph. Fischer’s exact and Mann-Whitney U tests were conducted for (G) and (H), respectively.

To confirm these results in a T-cell line, we performed drug TOA assays with RPV and DTG using the SupT1 T-cell line and VSV-G pseudotyped HIV-1 encoding an mScarlet reporter (**Fig. 5E and F**). Consistent with the findings described above (**Fig. 5A and Extended data 5)**, NC-G19S exhibited a statistically significantly shorter time lag between the completion of reverse transcription and integration than WT (∼2.4 h for NC-G19S vs. 5.5 h for WT) (**Fig. 5E and F**). Collectively, these results demonstrate that, in multiple target cell types and viral vector systems, the NC mutants exhibit a compression of the time between completion of reverse transcription and integration.

### NC-G19S does not alter the efficiency or kinetics of uncoating in the nucleus

Recent studies have demonstrated that intact viral cores enter the nucleus where reverse transcription is completed, and uncoating takes place near the site of integration ^19,68–71^. NC has been reported to bind and compact reverse-transcribed DNA within the core, likely playing a role in containing the internal pressure exerted by the newly synthesized viral DNA and preventing premature uncoating ^19,21^. To investigate the effect of the prototypical NC mutation NC-G19S on uncoating, we assessed uncoating efficiency in the nucleus using live-cell imaging of HIV-1 cores labeled with GFP-tagged CA in HeLa cells expressing mRuby-Lamin B ^19,68,69^. This analysis revealed no significant differences in the efficiency or kinetics of uncoating within the nucleus (**Fig. 5G and H**).

### HIV-1 acquires mutations in Env, NC and IN in the presence of DTG in primary T cells

Our previously reported DTG selection experiments were performed in immortalized T-cell lines ^49^. To gain further insights into INSTI resistance pathways under more physiologically relevant conditions, we conducted long-term propagation experiments with increasing DTG concentrations in peripheral blood mononuclear cells (PBMCs). We infected PBMCs with three different HIV-1 strains: the subtype B, X4-tropic, NL4-3 strain (**Fig. 6A**); the subtype B, R5-tropic, transmitted-founder isolate CH077 (**Fig. 6B**); and the subtype C, R5-tropic transmitted-founder strain CH185 (**Fig. 6C**). Over 6-12 months of continuous propagation, we were able to gradually increase the DTG concentration. To identify the genetic changes responsible for DTG resistance, we extracted genomic DNA from infected cells and performed sequencing analysis across the HIV-1 genome. Consistent with our previous study using the SupT1 T-cell line ^49^, the NL4-3 strain sequentially acquired multiple Env mutations and mutations in NC (NC-N27I) and IN (IN-S153Y and IN-D167H). Similarly, the CH077 molecular clone acquired three Env mutations (V208A, R577K, and D743N), NC-N27K, and IN-T122I. CH185 acquired and subsequently lost NC-A30T, and acquired two Env mutations (A533V and L764M), another NC mutation (NC-D48N), and two IN mutations (IN-E163K and R263K). We did not identify any mutations in the 3’PPT G-tract region^38–43^ in the viral strains tested here. Inactivating mutations in Vpu were also frequently detected, consistent with previous reports from us and others ^47,72,73^. These observations indicate that DTG resistance pathways shown here are independent of viral isolate and cell type. Notably, the identified NC mutations are within the zinc-finger domains. Previous studies suggest that the mutation at IN-T122 may contribute to a higher level of resistance to INSTIs when coupled with G140S/Q148H^74,75^. IN-S153Y, E163K, and R263K are reported as low-level DTG resistance mutations in the Stanford HIV Drug Resistance Database^76^.

**Fig. 6.**
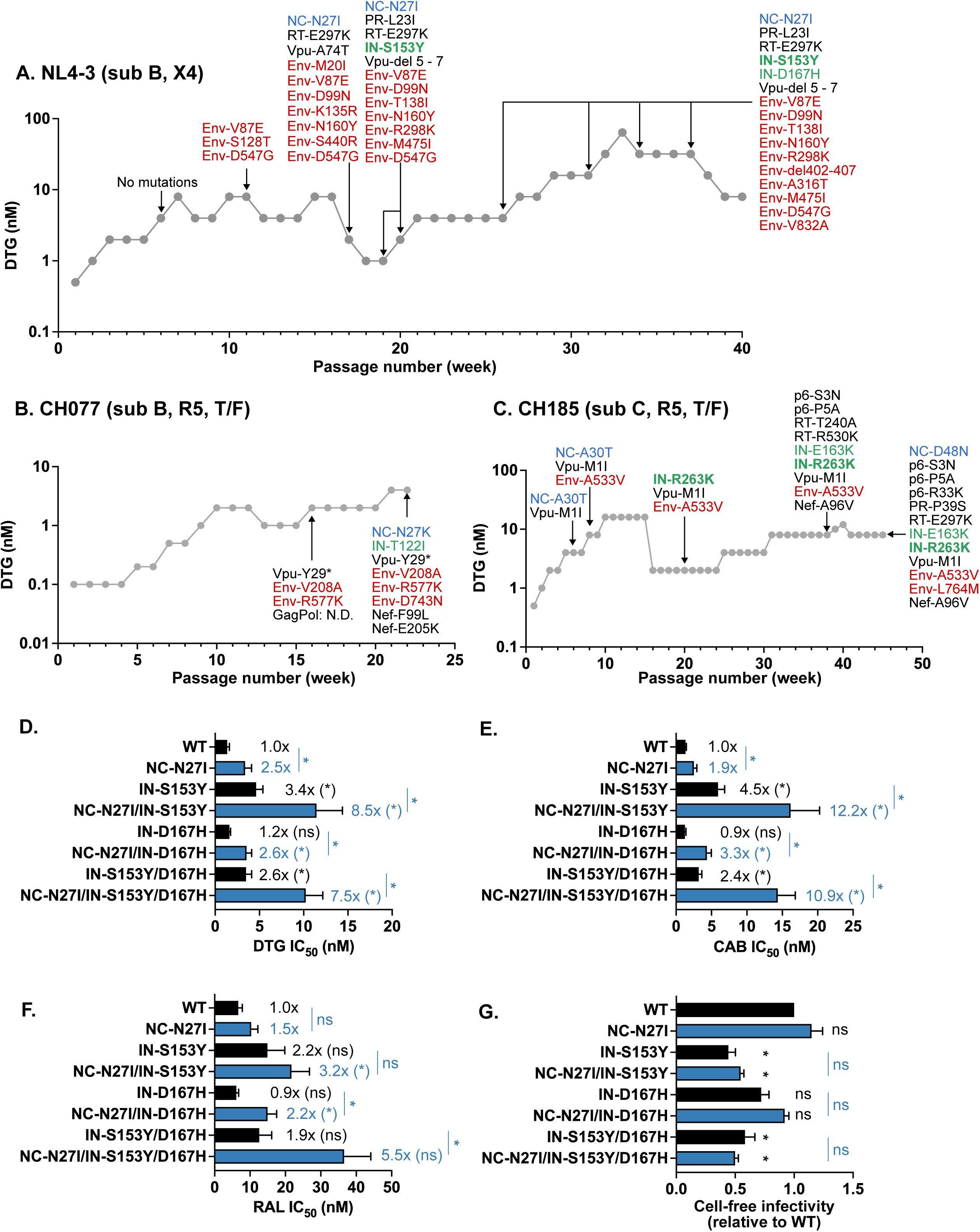
Selection of DTG-resistant variants in PBMCs. Long-term passaging of (A) NL4-3, (B) CH077, and (C) CH185 molecular clones in the presence of DTG. Activated PBMCs were infected with the indicated viruses to initiate the passaging experiments. At the time points indicated by the arrows, genomic DNA was extracted from the drug-treated cultures, and the Gag-Pol, and Env-Nef coding regions were sequenced. Mutations detected at frequencies greater than 25% in the bulk sequencing are shown. Mutations highlighted in bold are established DTG resistance mutations in IN. (D) DTG, (E) RAL, and (F) CAB sensitivity of the NL4-3 NC/IN variants identified during long-term passaging of NL4-3 in the presence of DTG (panel A). TZM-bl cells were incubated with RT-normalized viruses in the presence of various concentrations of the indicated INSTIs. (G) Cell-free infectivity of NL4-3 NC and IN variants in TZM-bl cells. Relative infectivity is shown, normalized to 1 for WT NL4-3. The data are shown as means ± SEM from >3 independent experiments. Statistical significance between variants with and without the NC mutation was determined using an unpaired t-test, with significance indicated as *p < 0.05. For panels D-G, (*) and (ns) indicate comparisons to the WT.

Interestingly, we observed in the PBMC selections that NC and IN mutations often arose together, suggesting that mutations in these two coding regions may work in concert to increase levels of INSTI resistance. To test this hypothesis, we introduced the NC/IN mutations identified in **Fig. 6A** into the NL4-3 molecular clone and assessed their effect on sensitivity to INSTIs (**Fig. 6D–F**). We found that IN-S153Y and IN-S153Y/D167H conferred 3.4-fold and 2.6-fold resistance to DTG, respectively. NC-N27I alone showed a statistically significant level of resistance to second-generation INSTIs DTG and cabotegravir (CAB), but not to the first-generation INSTI RAL. The level of resistance conferred by the NC-N27I mutation was similar to that of the IN-S153Y and IN-S153Y/D167H mutations. The combination of NC-N27I and IN-S153Y resulted in greater increases in DTG and CAB resistance (8.5-fold and 12.2-fold resistance, respectively) than mutations in either IN or NC alone. Although NC-N27I and IN-S153Y alone did not show substantial resistance to RAL, in combination they conferred a significant level of RAL resistance. While NC-N27I enhanced the resistance conferred by the co-selected IN mutations, this NC mutation did not rescue the ∼2-fold infectivity defect imposed by the IN mutations (**Fig. 6G**). These results demonstrate that NC and IN mutations that emerge together in PBMCs act in concert to increase INSTI resistance.

## Discussion

In this study, we characterized a panel of NC mutants selected in the presence of DTG. We demonstrated that INSTI resistance conferred by NC mutations is closely associated with accelerated early post-entry events in the HIV-1 replication cycle. Notably, while these NC mutations do not have a major effect on virus assembly or infectivity, they significantly shorten the time between completion of reverse transcription/uncoating and viral DNA integration. These results demonstrate that the NC mutations narrow the time window during which INSTIs can effectively inhibit integration, offering a novel mechanism for INSTI resistance.

Unlike canonical HIV-1 drug resistance mutations that directly alter drug-target interactions^32^, INSTI resistance mediated by NC mutations must involve alternative mechanisms, as INSTIs do not target NC. One possible scenario is that mutations in NC impair its chaperone activity during reverse transcription, leading to the formation of aberrant viral DNA ends that are not good substrates for INSTI binding. Point mutations in the CCHC motif of the NC zinc-finger domains have been shown to severely impair viral infectivity, at least partly due to incomplete viral DNA processing and formation of aberrant viral DNA ends^29^ (and confirmed here). In contrast, the NC mutations that arose in the presence of DTG ^49^ do not diminish particle infectivity or viral replication kinetics, and the integration efficiency of NC-G19S is higher than that of WT at early time points post-infection. These findings lead to the prediction that the NC mutations have a minimal effect on viral DNA ends. Indeed, using the NC-G19S mutant, we found that the sequence of 2-LTR junctions and virus-host junctions of integrated DNA are similar to those of WT. Furthermore, our competitive abrogation assay demonstrates that the NC mutations do not compromise functional intasome formation. Taken together, these observations indicate that DTG-selected NC mutations do not modify the INSTI-intasome interaction through alteration of viral DNA ends. Our work also demonstrates that resistance mediated by the selected NC mutations is mechanistically distinct from that conferred by 3’PPT mutations, as the latter are characterized by integration-independent viral protein expression and a bypass of INSTI inhibition ^38,40,41^. In addition, the 3’PPT mutant tested here did not replicate in an HTLV-1 Tax-negative cell line (the SupT1 cell line), whereas the NC mutants selected in the presence of DTG replicated with kinetics similar to those of the WT.

As mentioned in the Introduction, HIV-1 NC has been reported to possess the ability to condense viral DNA and stimulate integration^17,20–27^. A consistent finding across different drug TOA assay formats is that the NC mutants selected in the presence of DTG reduce the time between completion of reverse transcription or uncoating and viral DNA integration. Because INSTIs act by binding the intasome – a complex of newly synthesized and 3’ processed viral DNA ends and a multimer of IN ^5,7,8^ – reducing the time between viral DNA end formation and integration would effectively narrow the window of opportunity for INSTIs to bind the intasome and block integration.

HIV-1 preferentially integrates in SPADs – transcriptionally active, gene-dense regions of the host cell chromatin – and disfavors transcriptionally inactive regions of the chromatin, specifically lamina-associated domains (LADs)^64^. In addition to showing more viral DNA integration at early time points relative to the WT, NC-G19S also shows a subtle but statistically significant shift in integration out of SPADs and into LADs. This shift is much less pronounced than that previously reported for CA mutants unable to interact with the host protein CPSF6 or in cells depleted for CPSF6 ^68,77–79^. We speculate that NC may have some role in integration site selection, or the faster kinetics of NC-G19S integration may cause an increase in integration into LADs, which tend to be localized to the nuclear periphery. Regardless of the mechanism, these results underscore the need for further investigation into the timing of uncoating, intasome formation, and interactions with host factors during infection and the role of NC in these events.

Several INSTI resistance mutations in IN have been reported in both in vitro and clinical settings ^37,80–82^. Many of these IN mutations confer only a modest level of resistance to DTG^80^. For instance, a major DTG resistance mutation, IN-R263K, reduces DTG susceptibility by ∼3-fold ^80^. Notably, there is evidence that the emergence of mutants that exhibit only ∼4-fold resistance to DTG increases the risk of VF^83^. The level of resistance conferred by the NC mutations reported here is comparable to those conferred by these IN-resistance mutations, suggesting that the presence of NC mutations could contribute to VF in PWH. In our previous study ^49^ and in this current work, we have shown that NC mutations emerge along with Env and IN mutations in a T-cell line and in primary T cells. Previous work demonstrated that INSTI-resistance mutations in IN often decrease viral fitness^75,80,84^, but Env mutations rescue these defects ^47–49^. Notably, the NC mutations selected in the presence of DTG can increase the levels of INSTI resistance conferred by mutations in IN, indicating that NC mutations act in concert to reduce the susceptibility of HIV-1 to INSTIs.

In summary, this study provides new insights into the role of NC in the HIV-1 replication cycle and underscores the importance of mutations outside drug-target genes in the development of INSTI resistance.

## Material and Methods

### Cell lines and reagents

Human embryonic kidney (HEK) 293T cells, TZM-bl cells^85^, HeLa cells (ATCC CCL-2), HeLa:mRuby-LaminB cells [a HeLa-derived cell line constitutively expressing an mRuby-LaminB fusion protein^69^] were maintained in Dulbecco modified Eagle’s medium supplemented with 10% fetal bovine serum (FBS) at 37°C in 5% CO_2_. The SupT1 T-cell line and SupT1huR5 T-cell line^86^ were cultured in RPMI-1640 medium supplemented with 10% FBS at 37°C in 5% CO_2_. The SupT1huR5 T cell line was a gift from J. Hoxie (Perelman School of Medicine, University of Pennsylvania, Philadelphia, PA). DTG and BMS-378806 were purchased from MedChemExpress, and RPV and CAB from Cayman Chemical. The anti-HIV-1 Gag antibody (ab63917) was purchased from Abcam. LEN and anti-RT antibody (8C4) were obtained through the National Institutes of Health (NIH) HIV Reagent Program. Primary CD4 T cells were isolated from human peripheral blood mononuclear cells (PBMCs) using EasySep Human CD4+ T-cell isolation kit (STEMCELL Technologies) according to the manufacturer’s instructions. PBMCs were isolated using a ficoll gradient and stimulated with 2 μg/ml phytohemagglutinin (PHA-P) for 3 to 5 days before infection and then cultured in 50 U/ml interleukin-2 (IL-2).

### Plasmids

The full-length HIV-1 molecular clones pNL4-3^87^, pCH185^88^ and pCH077^89^ were used in this study. A series of pNL4-3 NC mutants were described previously^49^. pCH185 and pCH077 were kind gifts from C. Ochsenbauer and J. Kappes (University of Alabama at Birmingham). The pNL4-3/KFS clone, which does not express Env, was described previously^90^. pNL4-3 IN-D116N was described previously^50^. The NC regions with the indicated mutations were cloned into the pNL4-3/KFS using SpeI and SbfI restriction sites. NC-H23C and H44C ^29^ were introduced into pNL4-3 and pNL4-3/KFS by overlap PCR. The VSV-G–expressing plasmid, pHCMV-G, was a gift from J. Burns^91^. The NL4-3-derived mScarlet reporter virus plasmid, pNL4-3 mScarlet.6ATRi KFS and NL4-3 nanoLuc reporter virus plasmid, pNL4-3 nanoLuc.6ATRi KFS were constructed based on previously described pNL-LucR.6ATRi-BaL.ecto^51^. The genetic information and the related DNA materials were kind gifts from C. Ochsenbauer (University of Alabama at Birmingham). The DNA fragments containing the modified encephalomyocarditis virus (EMCV) 6ATR internal ribosome entry site (IRES) element (6ATRi) to replace the *renilla luciferase* with *mScarlet-I3* or *nanoLuc* were synthesized by Twist Bioscience. The DNA fragments were cloned into pNL4-3/KFS using HpaI and XhoI restriction sites. To increase the genetic stability of *mScarlet-I3* and *NanoLuc*, CG dinucleotides in the reporter-coding genes were removed after the codon optimization^52^. The HIV-1 helper construct pcHELP lacks both the packaging signal and primer-binding site and expresses all viral proteins except Nef and Env^92^. pcHELP-GFPCA is a derivative of pCHELP in which GFP is inserted between the MA and CA domains of Gag, producing a GFP-CA fusion protein following proteolytic processing during viral maturation ^19^. The NC-G19S mutant derivatives were generated by inserting a gBlock (Integrated DNA Technologies [IDT]) containing the G19S substitution in the Gag coding region, followed by subcloning into both pCHELP and pcHELP-GFPCA to yield pCHELP-G19S and pcHELP-GFPCA-NCG19S, respectively. The HIV-1-based vector pHmNG-ΔGag contains an mNeonGreen (mNG) reporter gene in place of the *nef* gene and lacks Env expression. A frameshift mutation was introduced into *gag* by SphI digestion followed by removal of 3’ overhang nucleotides using Klenow fragment, resulting in a premature stop codon.

### Preparation of virus stocks

HEK293T cells were transfected with HIV-1 proviral DNA using Lipofectamine 2000 (Invitrogen) according to the manufacturer’s instructions. VSV-G–pseudotyped viruses were prepared by cotransfecting HEK293T cells with pNL4-3/KFS and pHCMV-G at a DNA ratio of 10:1. At 48 h post-transfection (26 h for VSV-G-pseudotyped virus), virus-containing supernatants were filtered through a 0.45-μm membrane filter (Merck Millipore). The amount of virus in the supernatant was quantified by RT assay as previously described (Hikichi et al., Sci Adv, 2024). To generate GFP-CA-labeled virions containing the mNG reporter genome, 293T cells were co-transfected using polyethyleneimine (PEI; Polysciences) with the following plasmids: pCHELP-GFPCA and pCHELP (or their NC-G19S variants) at a 1:10 plasmid ratio (total of 3 µg DNA), pHmNG-ΔGag (7 µg), and pHCMV-G (2 µg). Approximately 24 h post-transfection, virus-containing supernatants were collected, filtered, and concentrated by ultracentrifugation (100,000 × g for 1.5 h at 4 °C) through a 20% (wt/vol) sucrose cushion in 1x phosphate buffered saline (PBS).

### Western blotting of viral proteins

Cell- and virus-associated proteins were solubilized in lysis buffer (30 mM NaCl, 50 mM Tris-HCl pH 7.5, 0.5% Triton X-100, 10 mM iodoacetamide, complete protease inhibitor [Roche]). Lysates boiled in 6× loading buffer (7 ml 0.5 M Tris-HCl/0.4% SDS, 3.8 g glycerol, 1 g SDS, 0.93 g DTT, 1.2 mg bromophenol blue) were subjected to SDS–polyacrylamide gel electrophoresis and transferred to polyvinylidene difluoride membranes (Merck Millipore). After blocking with Azure Fluorescent Blot Blocking Buffer (Azure Biosystems), membranes were probed with indicated antibodies at 4°C overnight, then incubated for 1 h with species-specific AzureSpectra Fluorescent Secondary Antibodies (Azure Biosystems). Bands were detected by fluorescence with a Sapphire Biomolecular Imager (Azure Biosystems) and quantified using ImageJ software.

### Quantification of viral RNA in the virion

Before viral RNA extraction, the virus supernatant from transfected 293T cells was treated with 20U of DNase I (ThermoFisher Scientific) at 37 °C for 1 h. Viral RNA was extracted with QIAamp Viral RNA Kits (Qiagen). Viral RNA was detected by QuantStudio 3 Real-Time PCR systems and TaqMan Fast Virus 1-Step Master Mix (ThermoFisher Scientific) using the following primer/probe set ^93^: forward primer GagF1 (5’-CTAGAACGATTCGCAGTTAATCCT-3’), reverse primer GagR1 (5’-CTATCCTTTGATGCACACAATAGAG-3’), and probe P-HUS-103 (5’-FAM-CATCAGAAGGCTGTAGACAAATACTGGGA-TAMRA-3’). The cycling conditions were: reverse transcription (50 °C, 5 min and 95 °C, 20 sec) and 40 cycles of denaturation (95 °C, 3 sec) and extension (60 °C, 30 s).

### Viral infection and DNA isolation

Before infection, VSV-G-pseudotyped viruses were treated with 20U of DNase I at 37 °C for 1 h. 5.0 x 10^5^ SupT1 T-cells were exposed to DNase I-treated, VSV-G-pseudotyped virus and placed at 4°C for 30 min for entry synchronization. Viral inputs were normalized by RT activity. Cells were incubated at 37 °C for 3, 6, 9, 12, and 24 h post-synchronization. At indicated times, total DNA was extracted using QIAamp DNA Blood Mini Kit (Qiagen), and viral DNA was measured by qPCR. As a negative control, VSV-G-pseudotyped viruses were heat-inactivated at 95 °C for 1 h.

### Sequence analysis of 2-LTR junction

24 h post-infection with VSV-G-pseudotyped viruses, total DNA from infected SupT1 T-cells was extracted as described above. The 2-LTR circle junction was amplified using PrimeSTAR GXL DNA polymerase (Takara Bio) with primers ^93^: forward primer SSF4 (5′-TGGTTAGACCAGATCTGAGCCT-3′) and reverse primer LTR-R5 (5′-AGGTAGCCTTGTGTGTGGTAGATCC-3′). The PCR amplicon was purified and cloned into pCR-Blunt II-TOPO vector (ThermoFisher Scientific) per the manufacturer’s instructions. The 2-LTR junction sequence was determined by Sanger sequencing (Poochon Scientific) using M13 reverse primer.

### Integration site analysis

HIV-1 integration site analysis (ISA) was performed as in ^94,95^. Briefly, 4 µg genomic DNA from each experimental condition was fragmented, A-tailed, and linker-ligated (NEBNext Ultra II FS, E7805L). Linker-mediated PCR targeted amplification and indexing of both 5’LTR and 3’LTR HIV-human integration junctions and the prepared library was sequenced by Illumina MiSeq with paired end reads (Illumina MS-102-2002). Canonical integration site analysis (complete LTR) and genome preference analysis was performed using a custom bioinformatic pipeline described in ^95^.

Aberrant integration site analysis (non-canonical LTR junctions) was performed using an adapted bioinformatic pipeline from ^63^. The fastq files were prepared by being converted to their reverse complements using Seqtk (https://github.com/lh3/seqtk) to accommodate the input requirement of the pipeline described in ^63^. 5’LTR and 3’LTR were analyzed individually using the pipeline because they were sequenced separately, and the results for the same sample were integrated afterward. Primers for ISA are the following: 3LTR_PCR1 (5’-TGTGACTCTGGTAACTAGAGATCCCTC-3’), 5LTR_PCR1 (5’-TCAGGGAAGTAGCCTTGTGTGTGGT-3’), TLinkerMID_P1 (5’-AGTTCAGACGTGTGCTCTTC-3’), 3LTR_PCR2 (5’-AATGATACGGCGACCACCGAGATCTACACAGGCGAAGACACTCTTTCCCTACACGACGCTCTTCCGATCTNNNNNNTCATCGAGCCTTTTAGTCAGTGTGGAAAATC-3’), 5LTR_PCR2 (5’-AATGATACGGCGACCACCGAGATCTACACATAGAGGCACACTCTTTCCCTACACGACGCTCTTCCGATCTNNNNNNCTACTATATGTCTTCTTTGGGACCAAATTAGCC-3’), Linker_P2* (5’-CAAGCAGAAGACGGCATACGAGATXXXXXXXXGTGACTGGAGTTCAGACGTGTGCT CTTCCGATC-3’). *XXXXXXXX is 8nt index for (de)multiplex of individual samples.

### Live-Cell Imaging

Live-cell imaging of HIV-1 uncoating was performed as previously described ^19^. Briefly, HeLa:mRuby-LaminB cells were seeded at 4 × 10^4^ cells/well in ibiTreated μ-slides (Ibidi) one day prior to infection. Cells were challenged with GFP-CA-labeled viruses at a low multiplicity of infection (MOI; ∼2 nuclear particles/cell) with polybrene (10 µg/ml; Sigma) via spinoculation (1,200 x *g* for 1 h at 15°C), allowing virus binding without internalization ^96^. Following spinoculation, the medium was replaced with prewarmed media to initiate viral internalization (defined as time 0). Aphidicolin (10 µg/ml) was added at the time of infection to inhibit cell division. Live-cell imaging was performed using a Nikon Eclipse Ti-E microscope equipped with a Yokogawa CSU-W1 spinning disk confocal unit, Plan-Apochromat 100x N.A. 1.49 oil immersion objective, and 488-nm (GFP) and 561-nm (mRuby) lasers. Images were acquired using a 561-nm long pass dichroic mirror and two ORCA-fusion BT cameras (Hamamatsu). A Tokai Hit stage-top incubator was used to maintain cells at 37°C. Time-lapse z-stacks (12 slices at 0.4 µm intervals) were collected every 20 minutes starting approximately 4 h post-infection and continued for 18 h. Nuclear GFP-CA spots were manually tracked using Nikon Elements, and their disappearance times were recorded.

### Quantification of viral cDNA in the infected cells

Viral and cellular DNA was quantified by qPCR using QuantStudio 3 Real-Time PCR systems and TaqMan Advanced Fast qPCR mix (ThermoFisher Scientific) with the following primer/probe sets ^93,97^: late RT products (U5Ψ: forward primer HIV LTR_GagF 5’-GAGATCCCTCAGACCCTTTTAGT-3’, reverse primer HIV LTR_GagR 5’-TTCAGCAAGCCGAGTCCT-3’, probe P-HUS-SS1 (5′-FAM-TAGTGTGTGCCCGTCTGTTGTGTGAC-TAMRA-3′), 2-LTR circle: forward primer SSF4 (5′-TGGTTAGACCAGATCTGAGCCT-3′), reverse primer LTR-R5 (5′-AGGTAGCCTTGTGTGTGGTAGATCC-3′), probe P-HUS-SS1. Host CCR5 was detected using forward primer CCR5-For (5′-CCAGAAGAGCTGAGACATCCG-3′), reverse primer CCR5-Rev (5′-GCCAAGCAGCTGAGAGGTTACT-3′), and probe P-CCR5-01 (5′-FAM-TCCCCTACAAGAAACTCTCCCCGG-TAMRA-3′). The cycling conditions were: initial denaturation (95 °C, 20 sec) and 40 cycles of denaturation (95 °C, 1 sec) and extension (60 °C, 20 sec). Standard curves for late RT product, 2-LTR, and CCR5 were based on pNL4-3, pJB1041 ^29^, and pRB2569 ^93^, respectively. To detect integrated DNA, the nested Alu-LTR PCR method was applied as previously described ^97^. The 1st PCR was performed using PrimeSTAR GXL DNA polymerase (Takara Bio) with Alu-HIV (5′-TCCCAGCTACTCGGGAGGCTGAGG-3′) and M661 (5′-CCTGCGTCGAGAGATCTCCTCTG-3′) primers. The 1st PCR cycling conditions were: initial denaturation (98 °C, 10 s), 22 cycles of denaturation (98 °C, 10 s), annealing (60°C, 15 sec), and extension (68 °C, 10 min). The 1st PCR product was diluted 10-fold and used as a template for measuring R/U5 DNA using M667 primer (5′-GGCTAACTAGGGAACCCACTGC-3′), AA55 primer (5′-CTGCTAGAGATTTTCCACACTGAC-3′), and HIV-FAM probe (5′-FAM-TAGTGTGTGCCCGTCTGTTGTGTGAC-TAMRA-3′).

### Single-round infectivity assay

Single-round infectivity assays were performed as previously described ^49^. TZM-bl cells (1.0 × 10^4^ cells) in 96-well plates were incubated with RT-normalized virus stocks. At the indicated time post-infection, luciferase activity was measured using the Britelite plus reporter gene assay system (PerkinElmer) and GloMax Navigator microplate luminometer (Promega). SupT1 or PHA-stimulated primary CD4+ T-cells (2.0 x 10^6^ cells) were incubated with the RT-normalized VSV-G pseudotyped nanoLuc reporter virus at 37°C for 2 h. Following incubation, the cells were washed and plated in 96-well flat-bottom plates and incubated at 37°C in the presence of various concentrations of DTG. Nano luciferase activity was measured using Nano-Glo luciferase system (Promega) according to the manufacturer’s instructions.

### Virus replication kinetics assays

Virus replication was monitored in SupT1 T cells as previously described ^49^. Briefly, SupT1 cells were incubated with the indicated pNL4-3 clones (1.0 μg of DNA/1.0 × 10^6^ cells) in the presence of DEAE-dextran (700 μg/ml) at 37°C for 15 min. Transfected cells (1.5 × 10^5^ cells) were plated in 96-well flat-bottom plates and incubated at 37°C in the presence of a range of concentrations of DTG. Aliquots of supernatants were collected to measure RT activity, and cells were split 1:3 every other day with fresh drug and medium. IC_50_ values were calculated on the basis of the AUC of the replication kinetics. IC_50_ was defined as the amount of inhibitor required to reduce the AUC by 50%.

### Time-of-drug-addition assay

TZM-bl cells (1.0 × 10^4^ cells) in 96-well plates were incubated with NL4-3 variants. SupT1 (5.0 x 10^5^ cells) or HeLa (4.0 x 10^4^ cells) in 48-well plates were incubated with VSV-G-pseudotyped NL4-3 mScarlet reporter virus at 4°C for 30 min for entry synchronization, followed by incubation at 37°C. The viral input was normalized by RT activity. At the indicated time post-infection, 3 µM BMS-378806 (for NL4-3 virus), 1 µM RPV, 100 nM LEN or 1 µM DTG was added. Luciferase activity was measured at 48 h post-infection using the Britelite plus reporter gene assay system (PerkinElmer) and GloMax Navigator microplate luminometer (Promega). mScarlet expression was measured at 48 h post-infection with an LSR-Fortessa (BD Biosciences) and analyzed by FlowJo software (Tree Star Inc.).

### Co-infection with VSV-G-pseudotyped GFP reporter virus and ‘dark virus’

The DTG abrogation assay was performed as previously described ^49^. The SupT1 T cell line was incubated with a fixed amount of VSV-G-pseudotyped GFP reporter virus with varying amounts of Env(-), VSV-G-pseudotyped HIV-1 (dark virus). Following incubation, the cells were washed and cultured in the presence or absence of drugs. At 48 h post-infection, GFP expression was monitored by flow cytometry using an LSR-Fortessa (BD Biosciences) and analyzed by FlowJo software (Tree Star Inc.).

### Long-term propagation of HIV-1 in PBMC in the presence of DTG

Selection of HIV-1 variants resistant to DTG was performed by serial passaging of the HIV-1-infected PBMC with increasing concentrations of DTG. Freshly thawed PBMCs were activated weekly with 50U/ml IL-2 and 2 µg/ml PHA-P. At 6-7 days post-infection, the infected cells were split and cultures were replenished with fresh drug and PBMCs. To monitor viral replication in the presence of DTG, a small portion of the infected cell culture was co-cultured with the SupT1huR5 T-cell line in a separate culture without DTG on day 5. Dose escalation was performed when cytopathic effects in the SupT1huR5 T-cell line were observed within a 2-day co-culture period. At the indicated time points, genomic DNA was extracted from infected cells using the DNeasy Blood and Tissue Minikit (Qiagen), and the Gag-Pol and Env-Nef coding regions were amplified by PrimeSTAR GXL DNA polymerase (Takara Bio) and sequenced by Sanger or Oxford Nanopore methods (Psomagen) using previously described primers ^98,99^.

## Ethics statement

PBMCs were obtained from anonymous, de-identified blood donors enrolled in the NIH Department of Transfusion Medicine Blood Products Program (NIH CC-DTM).

## Acknowledgements

Research in the Freed laboratory is supported by the Intramural Research Program of the Center for Cancer Research, National Cancer Institute, NIH. Y.H. was supported by a JSPS Research Fellowship for Japanese Biomedical and Behavioral Researchers at the NIH and an Intramural AIDS Research Fellowship from the Office of AIDS Research. We thank S. Hughes for discussions and J. Hoxie for providing the SupT1huR5 cell line. We are also grateful to C. Ochsenbauer and J. Kappes (University of Alabama, Birmingham) for providing the pCH185, pCH077, pNL-6ATRi.LucR constructs and technical information. The following reagents were obtained through the NIH HIV Reagent Program, NIAID, NIH: LEN and anti-RT antibody (8C4).

## Author contributions

Conceptualization: Y.H., E.O.F. Investigation: Y.H., R.C.B., S.C.P., S.D., E.C., B.L. Material and Software: Y.H., S.C.P., S.D., S.D.A., X.W. Supervision: X.W., V.K.P., E.O.F. Writing - original draft: Y.H., and E.O.F. Writing -review and editing: all authors

**Extended data 1. Replication kinetics of the NC mutants selected with DTG in the SupT1 T-cell line.** Replication kinetics of the indicated NL4-3 variants in the SupT1 T cell line in the absence or presence of DTG. Replication curves obtained in the presence of 0, 1, and 3 nM DTG are shown. Data are representative of three independent experiments.

**Extended data 2. NC mutations selected in the presence of INSTIs do not affect virion composition or viral release efficiency.** (A) The NC mutations investigated in this study. Mutated residues in the zinc-finger domain and basic linker are highlighted as red and blue, respectively. Control mutations in the CCHC motif are shown in green. Western blot analysis of the virus-producing 293T cells (B) and virion fraction produced from these cells (C), probed with anti-Gag Ab, anti-tubulin Ab, or anti-RT Ab. Representative western blots from at least three independent experiments are shown. Quantification of Gag processing in the virus-producing cells (D) and virions (E). Gag processing was expressed as the ratio of p24 compared to total p24 and Pr55Gag. (F) Virus release efficiency was expressed as the ratio of virion p24 compared to the total virion p24, cell p24, and cell Pr55Gag. The data are shown as means ± SEM from at least three independent experiments (G) Quantification of RT enzyme in the virion. The relative ratio of total RT (p66 and p51) per virion p24 is shown. The data are shown as means ± SEM from at least two independent experiments with statistical significance indicated (*p < 0.05) as per one sample t-test. (H) Viral RNA packaging in the virion. Viral RNA was extracted from virions and subjected to qRT-PCR. The viral RNA values were normalized by the quantification of total Gag bands (p24 and Pr55Gag) in the western blots shown in panel C. The data are shown as means ± SEM from at least two independent experiments with statistical significance indicated (*p < 0.05) as per one-sample t-test.

**Extended data 3. Analysis of 2-LTR circle junctions in total DNA isolated from infected SupT1 T-cells at 24 h post-infection.** PCR-amplified 2-LTR junctions were cloned into the pCR-Blunt II-TOPO vector for sequencing. (A) Representative examples of 2-LTR junctions. Junctions were categorized as having intact (GTAC) sequences, deletions (≥3 nucleotides), insertions, or small substitutions at the GTAC motif. "X" denotes any nucleotide, while "*" indicates deletions. Notably, extended U3 sequences with 3′PPT retention were observed exclusively in 2-LTR junctions of the 3′PPT-2C3A5T6Δ mutant. (B) The total number of sequences analyzed for each variant is indicated. n indicates the number of experimental repeats. (C) Statistical analysis of trends in 2-LTR modification. The chi-squared test was used to compare overall and individual trends in modified 2-LTR junction frequencies. The categories with significant differences were highlighted in light blue (*p < 0.05). ns: not significant. (E) Integration site frequency in cells infected with WT or NC-G19S. IS: integration site, RefSeq genes: genes annotated in the NCBI Reference Sequence Database, TSS: transcription start sites of RefSeq genes, SPAD: speckle-associated domains, LAD: lamina-associated domains. The binomial test was used to compare WT and NC-G19S (*p < 0.05). ns: not significant. (F) The frequency of the integration events with canonical and aberrant viral DNA ends at 5’LTR and 3’LTR. The binomial test was used to compare WT and NC-G19S (*p < 0.05). ns: not significant.

**Extended data 4. Cell-free infectivity and DTG sensitivity of NC-H44C and 3’PPT mutants.** (A) Cell-free infectivity of WT NL4-3, 3’PPT mutants, NC-H44C, and NC-G19S in TZM-bl cell. Relative infectivity is shown, normalized to 1 for WT NL4-3. The data are shown as means ± SEM from at least two independent experiments with statistical significance indicated (*p < 0.05) as per one sample t-test. (B) DTG sensitivity of the NC-G19S and 3’PPT mutants. TZM-bl cells were incubated with TCID_50_-normalized WT virus or the indicated mutants in the presence of a range of DTG concentrations. The data are shown as means ± SEM from >3 independent experiments.

**Extended data 5. Time-of-addition assay in HeLa cells using an HIV-1 reporter virus encoding mScarlet.** (A) HeLa cells were infected with VSV-G-pseudotyped mScarlet reporter virus, followed by the addition of RPV, LEN and DTG at the indicated time points. mScarlet expression at the indicated time points was normalized to the signals at 24 h post-infection. t_1/2_ for the indicated drugs is shown below the graph. The data are shown as means ± SEM from >3 independent experiments. (B) Differences in Δt_1/2_ (the difference in t_1/2_ between DTG and other drugs) between WT and the NC-G19S mutant.. Box plots show median, interquartile range, minimum, and maximum values. The mean value is indicated by a ‘+’. Statistical significance is shown (*p < 0.05) as determined by a one-sample t-test, ns: not significant.

