## Supplementary figures and images for "Elucidating the Mechanism by Which HIV-1 Nucleocapsid Mutations Confer Resistance to Integrase Strand Transfer Inhibitors"

### Extended data 1

**Extended data 1**

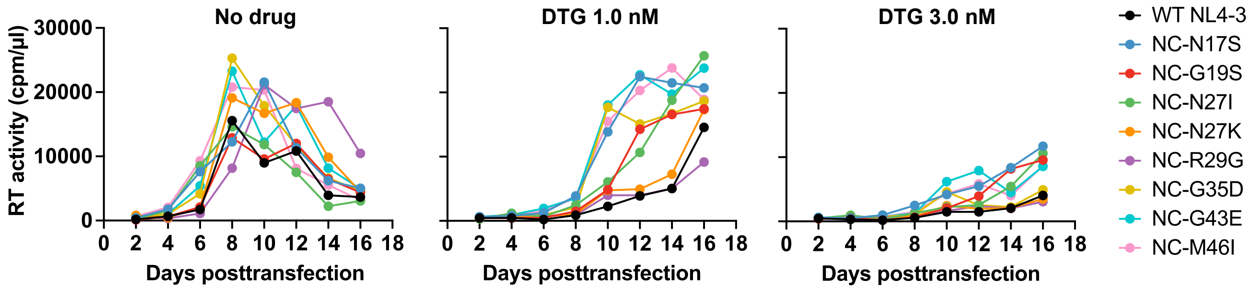

### Extended data 2

**Extended data 2**

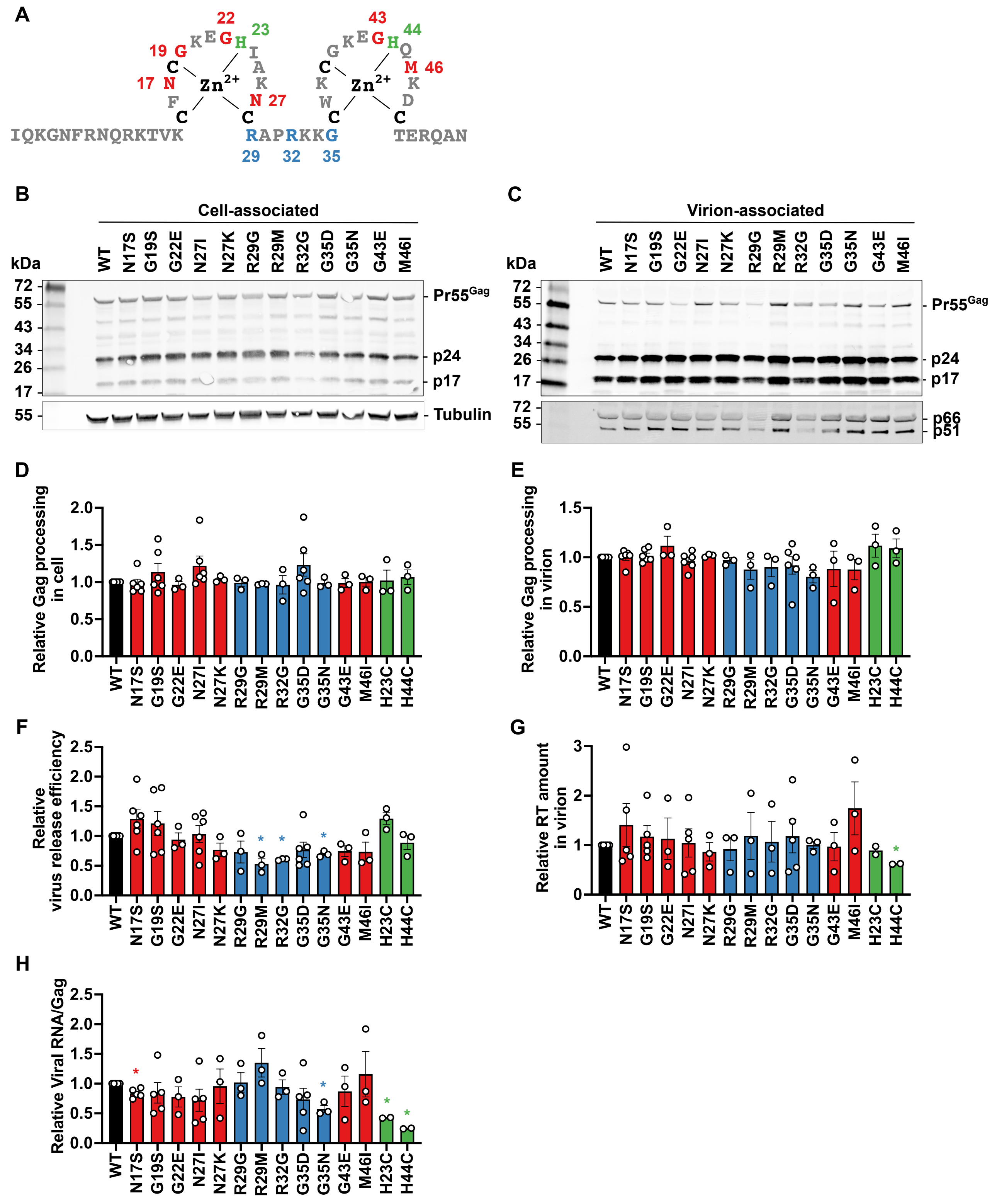

### Extended data 4

**Extended data 4**

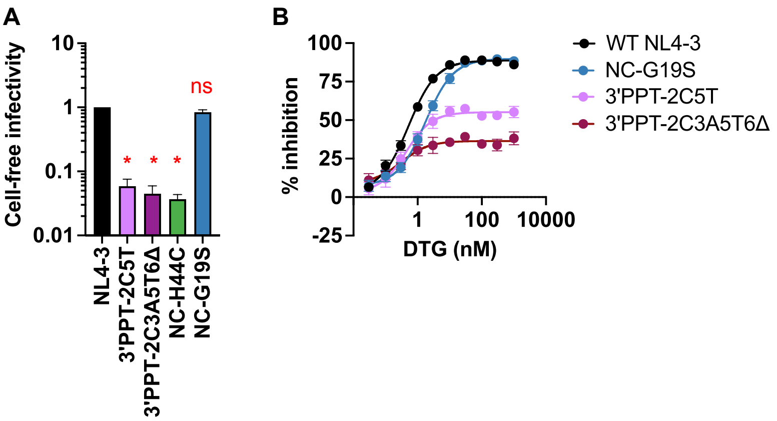

### Extended data 5

#### Extended data 5

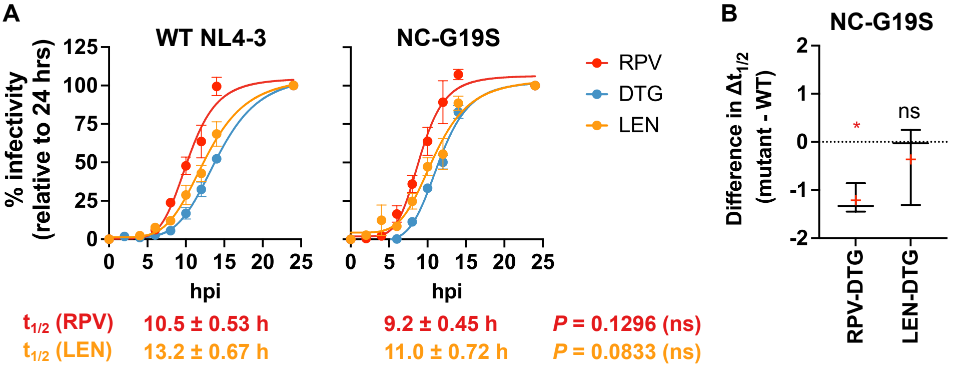


| | WT NL4-3 | NC-G19S | |
|---|---|---|---|
| $t_{1/2}$ (RPV) | 10.5 ± 0.53 h | 9.2 ± 0.45 h | *P* = 0.1296 (ns) |
| $t_{1/2}$ (LEN) | 13.2 ± 0.67 h | 11.0 ± 0.72 h | *P* = 0.0833 (ns) |
| $t_{1/2}$ (DTG) | 14.1 ± 0.13 h | 11.6 ± 0.23 h | *P* = 0.0007 (*) |
