## Extended data 3 for "Elucidating the Mechanism by Which HIV-1 Nucleocapsid Mutations Confer Resistance to Integrase Strand Transfer Inhibitors"

**A**

| | U5 | U3 | |
|---|---|---|---|
| | AGTGAATTAGCCCTTCCAGT | ACGCTAGAGATTTTCCAC | Consensus GTAC |
| | AGTGAATTAGCCCTTCCAXX--insert--XXGCTAGAGATTTTCCAC | | Insertion |
| | AGTGAA*****del******--insert--*****del*****TCCAC | | Insertion |
| | AGTGAA*****del****** | ****del****TTTCCAC | Deletion |
| | AGTGAA*****del****** | ACGCTAGAGATTTTCCAC | Deletion |
| | AGTGAATTAGCCCTTCCAGT | ****del****TTTCCAC | Deletion |
| | AGTGAATTAGCCCTTCCAXX | XXGCTAGAGATTTTCCAC | Substitution |
| | AGTGAATTAGCCCTTCCAGT | ACTGCTTTTCTTTTAAAA | Extended U3 with 3'PPT |

3'PPT retention

**B**

| | No. clones (n) |
|---|---|
| WT NL4-3 | 433 (5) |
| NC-G19S | 376 (4) |
| NC-N17S | 184 (2) |
| NC-N27I | 177 (2) |
| NC-H44C | 133 (2) |
| IN-D116N | 284 (3) |
| NC-G19S/IN-D116N | 287 (3) |
| NC-N17S/IN-D116N | 193 (2) |
| NC-N27I/IN-D116N | 189 (2) |
| NC-H44C/IN-D116N | 174 (2) |
| 3'PPT-2C3A5T6Δ | 92 (1) |

**C**

| | | *P* value | | | | | |
|---|---|---|---|---|---|---|---|
| | | Overall | Consensus | Insertion | Deletion | Substitution | Extended U3 |
| IN-WT | NC-G19S | 0.193 | 0.710 | 0.069 | 0.358 | 0.797 | N.D. |
| | NC-N17S | 0.145 | 0.155 | 0.842 | 0.145 | 0.750 | N.D. |
| | NC-N27I | 0.169 | 0.276 | 0.803 | 0.093 | 1.000 | N.D. |
| | NC-H44C | < 0.01 | 0.047 | < 0.01 | 0.734 | 0.012 | N.D. |
| | 3'PPT-2C3A5T6Δ | < 0.01 | 0.023 | < 0.01 | 0.126 | 0.418 | < 0.01 |
| IN-D116N | NC-G19S/IN-D116N | 0.705 | 0.393 | 0.633 | 1.000 | 0.746 | N.D. |
| | NC-N17S/IN-D116N | 0.181 | 0.087 | 0.086 | 1.000 | 1.000 | N.D. |
| | NC-N27I/IN-D116N | 0.226 | 0.087 | 0.268 | 1.000 | 0.549 | N.D. |
| | NC-H44C/IN-D116N | < 0.01 | < 0.01 | 0.256 | < 0.01 | 0.949 | N.D. |
| | 3'PPT-2C3A5T6Δ | < 0.01 | < 0.01 | 0.086 | < 0.01 | 0.478 | < 0.01 |

**D**

| | | WT | G19S | Binominal test *P* value |
|---|---|---|---|---|
| Total unique IS | | 46,758 | 44,265 | |
| In RefSeq genes (%) | | 72.35 | 71.85 | 0.0095 |
| In CpG islands (%) | +/- 1kb | 0.601 | 0.549 | 0.0059 |
| | +/- 2kb | 1.722 | 1.450 | < 0.001 |
| | +/- 5kb | 5.355 | 4.810 | < 0.001 |
| In TSS of RefSeq genes (%) | +/- 1kb | 0.768 | 0.834 | 0.0827 |
| | +/- 2kb | 2.318 | 2.465 | 0.0268 |
| | +/- 5kb | 10.231 | 9.768 | <0.001 |
| Median gene density in 1Mb region around IS | | 13 | 11 | 0.4270 |
| In LADs (%) | | 11.8 | 13.3 | < 0.001 |
| In SPADs (%) | | 23.5 | 20.2 | < 0.001 |

**E**

| | WT | G19S | Binominal test *P* value |
|---|---|---|---|
| Total counts | 50,366 | 48,079 | |
| Canonical ends | 46,758 | 44,265 | |
| Aberrant ends | 3,608 | 3,814 | < 0.001 |
| 3'LTR | 3,382 | 3,541 | < 0.001 |
| Deletions | 2,596 | 2,691 | < 0.001 |
| Insertions | 786 | 850 | < 0.001 |
| 5'LTR | 226 | 273 | < 0.001 |
| Deletions | 1 | 3 | 0.0721 |
| Insertions | 225 | 270 | < 0.001 |
| % canonical ends | 92.84 | 92.07 | |
| % aberrant ends | 7.16 | 7.93 | |
| 3'LTR | 6.71 | 7.36 | |
| 5'LTR | 0.45 | 0.57 | |
